# LOCATE: using Long-read to Characterize All Transposable Elements

**DOI:** 10.1101/2025.02.26.640385

**Authors:** Zhongren Hu, Bo Xu, Xiaoyan Zhang, Xiaou-ou Zhang, Zhiping Weng, Tianxiong Yu

## Abstract

Transposons constitute ∼45% of the human genome, driving gene evolution and contributing to disease, but their repetitive nature complicates the identification of new insertions. We present LOCATE (<u>Lo</u>ng-read to <u>C</u>haracterize <u>A</u>ll <u>T</u>ransposable <u>E</u>lements), an algorithm using long-read sequencing to detect and assemble transposon insertions. LOCATE outperforms existing tools on simulated datasets and achieves the best performance in two previous benchmarks, as well as in a new benchmark we constructed using real biological datasets. Applying LOCATE to public datasets revealed that pre-existing Alu copies create two hotspots for Alu and LINE1 insertions: the A-rich linker and the poly(A) tail. We further observed a preference for self-insertions over non-self-insertions in Alu and LINE1, suggesting a “feedforward” transposition mechanism in which Alu and LINE1 RNA transcripts target the hotspots of their source copies to generate new insertions. LOCATE enhances our ability to study transposons and their role in genome dynamics.

## Introduction

Transposons, or transposable elements, are DNA sequences capable of moving within the genome. They make up as much as 85% of the metazoan genomes and approximately 45% of the human genome (1, 2). While their activity can disrupt genome stability and lead to diseases like cancer (2–6), transposons also play essential roles in promoting genetic diversity and driving genome evolution (7–9). This dual nature makes the study of transposon movement crucial for understanding both its evolutionary significance and its impact on health. However, accurately characterizing transposon insertion events has remained a technical challenge due to their repetitive nature.

Many algorithms have been developed to identify new germline transposon insertions, including TEMP2, MELT, TELR, MEHunter, PALMER, TLDR, TrEMOLO, GraffiTE, and xTea (10–18). While some of these algorithms (i.e. TEMP2 and MELT) rely on short-read whole genome sequencing (DNA-seq), they struggle to detect transposon insertions in repetitive regions due to their inherent read-length limitations (10, 11). Long-read sequencing creates opportunities to address this limitation, but it brings technical challenges, such as higher error rates compared to short-read sequencing (19–22). Some existing algorithms detect transposon insertions by identifying reads partially aligned to both the genome and a transposon (12, 15, 16, 18), while others leverage genome assembly or structural variation analysis to specifically pinpoint transposon insertions (13, 14, 17). However, the high error rates in some long-read sequencing platforms can compromise alignment and assembly accuracy, particularly in repetitive regions. Even in platforms with low sequencing error rate (e.g. Pacbio high-fidelity sequencing), the complexity of the genome makes it challenging to distinguish real transposon insertions from other structure variations in repetitive regions.

All existing algorithms face one or more limitations, including species-specific applicability, inconsistent performance across sequencing platforms, errors in assembling insertion sequences, and, most importantly, failure to accurately detect “self-insertions” (e.g., an Alu element inserted within an pre-existing Alu element in the human genome) (10–18). These challenges highlight the need for an algorithm capable of accurately detecting transposon insertions across all genomic regions, assembling their sequences with minimal error, and being universally applicable to diverse species and long-read sequencing platforms.

## Materials and methods

### The LOCATE algorithm

The algorithm consists of four modules designed to ensure accuracy while maximizing sensitivity in detecting transposon insertions. The process begins with identifying candidate transposon insertions by aligning long reads to a reference genome using minimap2 (23). Parameters are tailored to the sequencing platform employed, such as PacBio, HiFi, or Oxford Nanopore Technologies, to ensure optimal alignment accuracy ("-aY --MD -x map-pb/map-hifi/map-ont"). Reads containing insertions or clipped segments longer than 100 base pairs are grouped into clusters if their breakpoints are within 50 base pairs. The inserted or clipped sequences from these clusters are subsequently aligned to transposon consensus sequences. Clusters where more than 80% of the reads align to transposons are designated as candidate insertions.

To minimize false positives caused by the repetitive nature of transposons, a machine learning framework removes alignment artifacts. This step relies on an AutoML model trained on simulated datasets generated specifically for this purpose. For larger genomes like the human genome, 150 datasets are simulated, whereas smaller genomes, such as *D. melanogaster*, require up to 300 datasets to ensure comprehensive training. The algorithm is applied to these simulated datasets, and false-positive insertions recurrently detected across datasets (in >10 human datasets or in >20 *D. melanogaster* datasets) are flagged as blacklists. These blacklists, which commonly occur near assembly gaps, are excluded from subsequent model training to prevent bias. Nineteen features of candidate transposon insertions are used in model training, including characteristics such as sequencing coverage, read length, genomic context, alignment quality, and the proportion of reads clipped at both ends (detailed in Table S7). The AutoML model is trained on 90% of the simulated datasets via AutoGluon TabularPredictor (version 1.0.0), with parameters “presets=best_quality” (24). Briefly, AutoGluon trains seven base machine learning models (LightGBM, CatBoost, XGBoost, RandomForest, ExtraTrees, KNearestNeighbors, and NeuralNetwork) and uses a multi-layer stacking strategy to ensemble predictions from each base model with the original features to make the final prediction. This model enables robust and accurate prediction of alignment artifacts. Tested on the remaining 10% of datasets, the AutoML models achieve 99.87–99.92% sensitivity and 99.88–99.93% precision in detecting human transposon insertions, and 99.00–99.36% sensitivity and 99.77–99.86% precision in detecting *D. melanogaster* insertions.

The third module involves assembling and characterizing the sequences of identified insertions. Supporting reads for each insertion are assembled with Wtdbg2 (v2.5) using the commands “wtdbg2 -l 256 -e 3 -S 1 –rescue-low-cov-edges –node-len 256 –ctg-min-length 256 –ctg-min-nodes 1” and “wtpoa-cns -c 1” (25). The resulting draft assemblies undergo polishing to correct alignment and mismatch errors, with the first round using Samtools consensus to improve overall sequence accuracy with parameters “--ff 3332 -m simple -c 0 -d 1-H 0.9” (26). To address homopolymer-length errors frequently observed in assemblies derived from long-read sequencing datasets, a second polishing step is performed. Supporting reads are aligned back to the Samtools-polished assemblies via minimap2, and the most common length for each homopolymer is determined. Reads with correct lengths are then used to generate consensus sequences for each homopolymer. Finally, all homopolymers are combined to generate the final polished transposon insertion sequences. These insertions are then characterized to annotate features such as poly(A) tails, truncations at the 5′ or 3′ ends, internal deletions, target site duplications, inversions, recombinations, and translocations In the final module, post-filtering refines the list of detected insertions by removing residual artifacts and ensuring biologically meaningful candidates are retained. Filtering criteria vary according to the transposon class and its integration mechanism. For retrotransposons without long terminal repeats, such as LINEs and SINEs, insertions lacking complete 3′ ends or poly(A) tails are excluded unless they are located in non-repetitive genomic regions and exhibit full-length characteristics. Insertions of LTR retrotransposons require the presence of complete 5′ and 3′ ends and, in repeat regions, validated target site duplications. Similarly, DNA transposons are retained if they possess complete termini and valid target site duplications when integrated into repetitive regions. These rigorous post-filtering steps enhance specificity and ensure the final set of detected insertions is biologically accurate and suitable for downstream analyses. LOCATE is freely available at Github: https://github.com/red-t/LOCATE.

### Data simulation

To benchmark LOCATE and other algorithms, we generated simulation datasets for human and *D. melanogaster* using SimulaTE and custom scripts (27). For the human dataset, we first generated 2,500 transposons, comprising 2,086 Alu, 301 LINE1, 97 SVA, and 16 HERV-K elements. Of these, 50% of the LINE1 elements were 5′ truncated, with truncation lengths ranging from 607 bp to 1,821 bp. These transposons were inserted into the hg38 reference genome, with 90% having frequencies of either 0.5 (heterozygous) or 1 (homozygous). The remaining 10% were simulated as mosaic insertions with frequencies ranging from 0.1 to 0.9. Using the read_pool-seq_pacbio.py module from SimulaTE, we simulated long reads for three platforms: PacBio HiFi, PacBio CLR, and ONT. The average error rates for HiFi, CLR, and ONT reads were 1%, 10%, and 7%, respectively, while their average lengths were 13.5 kbp, 7.9 kbp, and 7.2 kbp. Additionally, we simulated 150 bp paired-end short reads with an error rate of 0.05%.

Similarly, for *D. melanogaster*, we generated 500 transposon insertions representing 125 transposon subfamilies. We simulated 10% 5′ truncations for LINE elements and 10% internal truncations for LTR and DNA transposons. These transposons were inserted into the dm6 genome with frequencies ranging from 0.1 to 1. Long reads and short reads were simulated using the same strategies as for human.

To train the AutoML models for filtering alignment-induced artifacts, we simulated additional datasets for human and *D. melanogaster*. In total, 150 simulation datasets were generated for human and 300 for *D. melanogaster*. 90% of these datasets were used for model training, with the remaining 10% reserved for testing.

### Comparing LOCATE to other transposon insertion detection algorithms

LOCATE were benchmarked with nine additional transposon insertion detection algorithms: TEMP2, MELT, TELR, MEHunter, PALMER, TLDR, TrEMOLO, GraffiTE, and xTea (10–18).

These algorithms were evaluated using both simulated datasets and the GIAB HG002 datasets. We first downloaded human transposon consensus sequences from the MELT package (LINE1, Alu, and SVA) and Repbase (HERV-K) (11, 28). To annotate human transposon insertions in the reference genome, RepeatMasker was used with parameters “-s -no_is -norna -nolow -pa 32 -e ncbi -cutoff 255 -div 40 -frag 20000.” For *D. melanogaster*, transposon consensus sequences were sourced from FlyBase, and transposon insertion annotations were directly downloaded from RepeatMasker (29). All algorithms were executed following the best practices recommended on their respective websites. Since xTea and PALMER are specifically designed for human genomes, they were excluded from the benchmark analysis in *D. melanogaster*.

Many of these algorithms output both germline transposon insertions and de novo transposon insertions, the latter supported by a limited number of reads (10, 13, 15). To ensure a fair comparison, we applied the same read support thresholds for germline transposon insertions: a minimum of two supporting reads for datasets with 1–5x coverage, three reads for 10x dataset, and four reads for datasets with 20–50x coverage.

### Constructing a benchmark from GIAB HG002 datasets

The gold standard for transposon insertions in the GIAB HG002 dataset was established using multiple lines of evidence. First, insertion sequences were retrieved from LOCATE’s candidate insertions. If no sequence was available from LOCATE, sequence assembled by other algorithms or one of the supporting reads was used. Minimap2 (-k11 -w5 --sr -O4,8 -n2 -m20 --secondary=no) was employed to map insertion sequences back to the transposon consensus sequence for feature annotation. For sequences that failed to align with minimap2, BLAST (-best_hit_score_edge 0.1 -evalue 1) was used as an alternative. Insertions lacking sequences were excluded due to insufficient evidence of insertion signals

Next, insertions detected by more than five algorithms (1,421 in total) were directly included in the gold standard as the most reliable insertions. Additional validation was conducted using IGV [60] to exclude false positives among 792 candidates detected by fewer than five algorithms, which were required to exhibit complete 3′ termini.

Manual curation excluded 56 insertions with excessive coverage (Fig. S5A; >5 copy number), indicating mapping errors; 358 without generic transposon structure (Fig. S5B; 5′ and 3′ termini >500 bp from sequence ends or multiple transposon fragments with high proportion of non-transposon sequences); and 67 insertions primarily supported by multi-mapped reads, frequently accompanied by reads from >2 distinct haplotypes, suggesting mapping errors (Fig. S5C). A total of 311 insertions were validated and incorporated into the gold standard.

In summary, the GIAB HG002 gold standard dataset comprises 1,421 transposon insertions detected by more than five algorithms and 311 additional insertions validated through manual inspection.

### Preference of Alu and LINE1 insertion sites at pre-existing transposon copies

Transposon insertions detected by LOCATE, xTea, TLDR, TrEMOLO, GraffiTE, and PALMER from the GIAB HG002 HiFi dataset, and those detected by LOCATE from the 51 HPRC HiFi datasets were analyzed to assess their site preferences. Specifically, we calculated the enrichment of Alu and LINE1 insertions in three categories: pre-existing Alu copies, pre-existing LINE1 copies, and other regions. Enrichment was defined as the percentage of insertions in a specific region divided by the percentage that region represents across the reference genome. For insertions into Alu elements, enrichment was further calculated at each base pair of the Alu consensus sequence. The background for this analysis was defined as the percentage of each consensus nucleotide in the reference genome. LINE1 is much longer than Alu, and the number of insertions is not enough to assess insertion enrichment at base pair resolution.

### Speed of LOCATE and other algorithms

We benchmarked the speed of LOCATE alongside other transposon insertion detection algorithms on our high-performance cluster. Each algorithm was allocated 128 Gb of memory and 32 CPUs. The total runtime for each algorithm was recorded and compared to assess their computational efficiency.

### Statistical analysis

Statistics were computed with R (version 4.4.2) using Chi-squared test. Statistic methods, sample size and *p*-values can be found in figures, figure legends, or results. Significance was defined as *p*-value < 0.05. *: *p* < 0.05, ***: *p* < 0.001, n.s.: not significant.

## Results

### Design and implementation of LOCATE

We developed LOCATE (<u>Lo</u>ng-read to <u>C</u>haracterize <u>A</u>ll <u>T</u>ransposable <u>E</u>lements), an algorithm designed to accurately identify new germline transposon insertions and assemble their sequences. LOCATE operates through four sequential modules, each of the latter three modules addressing specific technical challenges (Fig. 1; see Methods). The first module extracts reads spanning or partially covering transposon insertions (supporting reads) and identifies candidate insertions using read clusters, similar to existing approaches.

**Figure 1.**
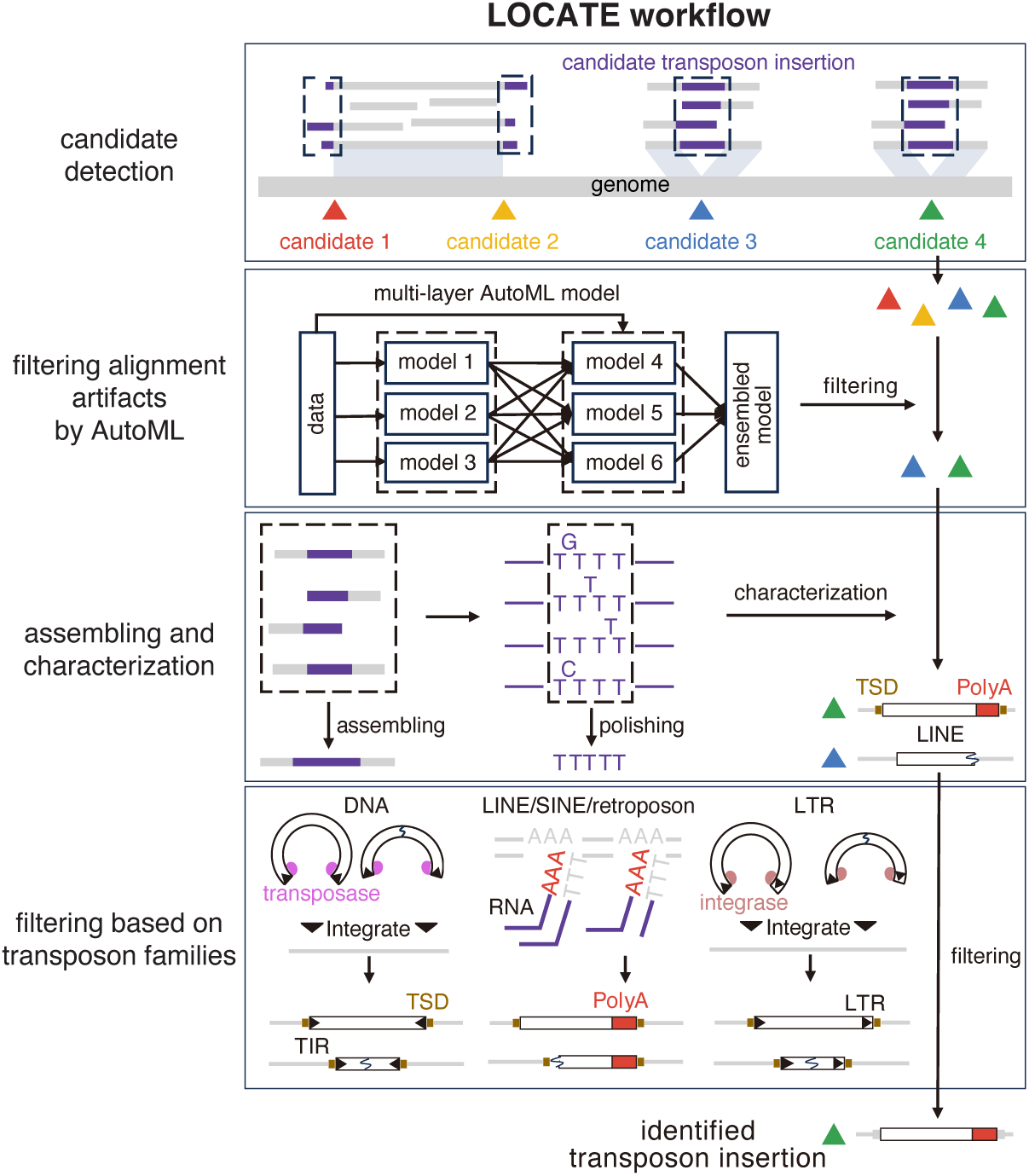
Overview of the LOCATE workflow. LOCATE consists of four modules. The first module identifies candidate transposon insertions by mapping reads to both the genome and transposon sequences. The second module applies machine learning models to filter out false positives resulting from inaccurate alignments. The third module assembles insertion sequences and annotates them. The final module filters false positives from other sources based on the characteristics of the identified insertions.

Imperfect read alignment, particularly in repetitive regions, introduces many false positive transposon insertions. The second module of LOCATE employs an automated machine-learning (AutoML) framework to filter out these artifacts. Using simulated long-read datasets with varying error rates and carefully selected alignment features—such as supporting read count, alignment quality, number of reads from 3′ and 5′ ends, and genomic context (e.g., repetitive elements or assembly gaps)—the AutoML model achieves exceptional performance in identifying artifacts induced by inaccurate alignments, with 99.9% sensitivity and precision in simulated human datasets and 99.2% sensitivity with 99.8% precision in simulated *D. melanogaster* datasets (see Methods).

A major advantage of long-read sequencing is its ability to assemble insertion sequences, revealing insertion characteristics such as divergence from consensus, target site duplications (TSDs), truncations, poly(A) tails, inversions, translocations, and recombinations. However, homopolymer-length errors in long reads pose a challenge for insertion sequence assembling (30, 31). Polishing via short-read sequencing is a common way to remove these errors but it is unreliable for repetitive elements, where short-reads cannot be unambiguously aligned (19, 32, 33). In order to assemble transposon insertions with minimal errors, the third module of LOCATE overcomes these errors by performing local assembly and polishing with Wtdbg2 and Samtools (25, 26), followed by a second polishing step that carefully inspects and curates the length of homopolymers. This module ensures high-quality assembly and structural characterization of transposon insertions.

Finally, LOCATE uses the characteristics of transposon insertions to filter artifacts derived from other sources, such as structure variations (e.g., duplication) involving pre-existing transposon insertions and imperfect reference genome assembly. LOCATE applies tailored post-filtering based on transposon families. LINEs, SINEs, and retroposons are identified by their characteristic poly(A) tails and complete 3′ ends, resulting from their target-primed reverse transcription (TPRT) mechanism, which starts with nicking of the transposon 3′ end to genome and can be prematurely terminated, resulting in truncated 5′ ends (34, 35). LTR transposons, integrated with the help of integrase binding to long terminal repeat (LTR) sequences, require complete 5′ and 3′ ends (36–40). As recombination between the two LTRs of a LTR transposon insertion occurs frequently, solo-LTR insertions are also allowed (41, 42). DNA transposons, recognized by transposase binding to terminal inverted repeats (TIRs) for genome integration, also need complete 5′ and 3′ ends (43, 44).

By integrating these four modules, LOCATE accurately detects transposon insertions, assembles their sequences, and characterizes their features across entire genomes, even in highly repetitive regions.

### Identification of transposon insertions in simulated human and *D. melanogaster* datasets

To evaluate the performance of LOCATE, we first compared it against other transposon insertion detection algorithms using simulated human datasets. We randomly put transposon insertions into the human reference genome (hg38) and simulated long-reads from this new genome in three sequencing platforms: PacBio high-fidelity (HiFi) sequencing, PacBio continuous long-read (CLR) sequencing, and Oxford Nanopore Technologies (ONT) sequencing (Figure S1; see Methods). Based on data from Genome In A Bottle (GIAB) and other studies (45, 46), different error rates and read lengths were simulated for each platform, ranging from 1% to 10% (Fig. S1; Table S1; see Methods). We also simulated datasets with varying coverages, to assess their effect on the algorithm performances. Additionally, we simulated short-read datasets for comparison with long-read-based algorithms. The simulated datasets represent the simplest scenario where artifacts are only derived from imperfect alignments: 1. the new genome used for data simulation only contains new transposon insertions, excluding the effects from other structure variations; 2. all the simulated reads are from the new genome, excluding the effects from unassembled contigs and other sources. LOCATE consistently outperforms other methods in detecting transposon insertions in simulated human datasets, achieving the highest sensitivity and precision across all platforms (Fig. 2A–C; Table S2).

**Figure 2.**
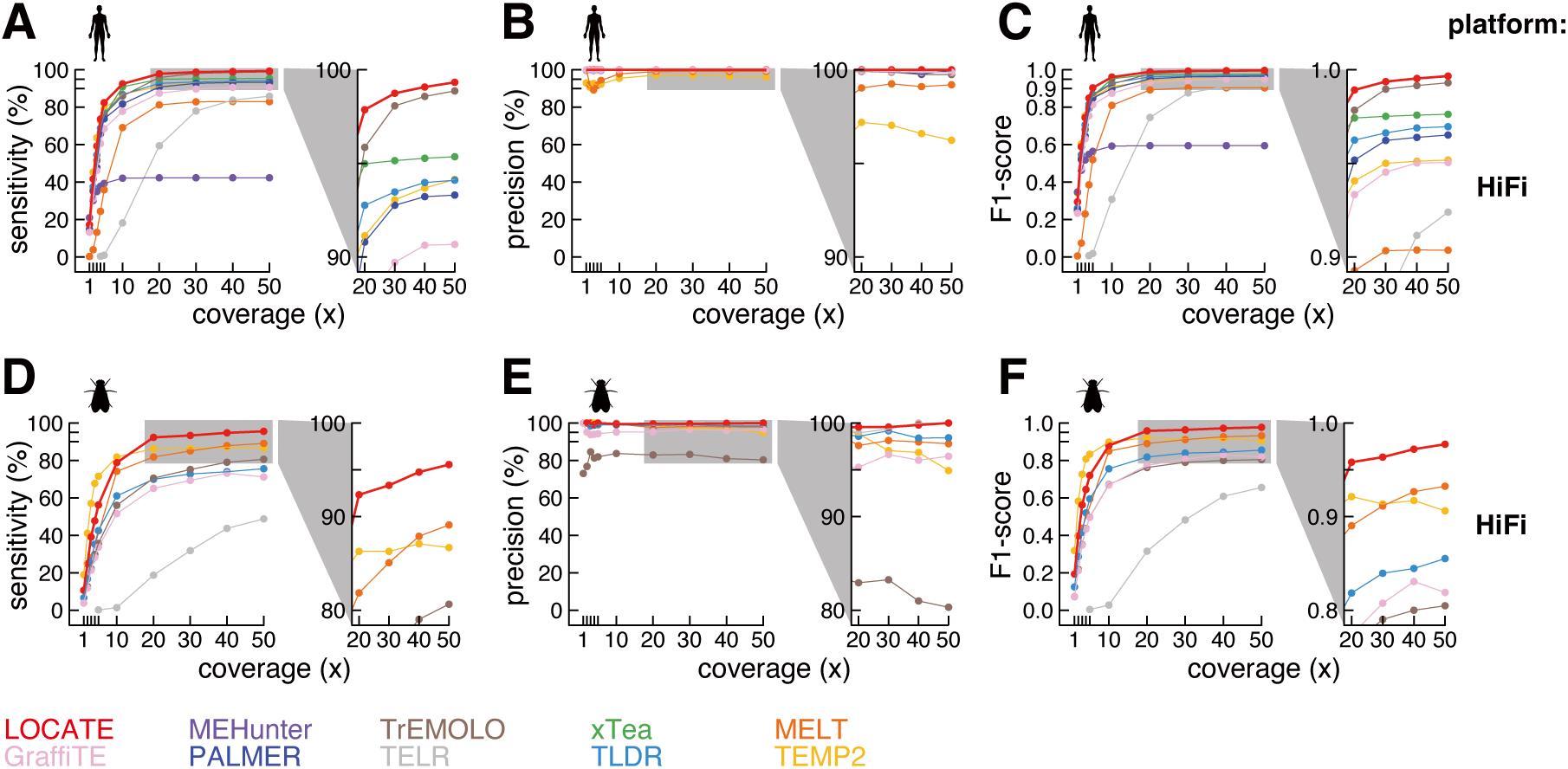
LOCATE outperforms other algorithms in detecting transposon insertions in simulated human and *D. melanogaster* datasets. The top figures show **A.** sensitivity, **B.** precision, and **C.** F1 score for each algorithm in detecting germline transposon insertions in simulated human Pacbio HiFi datasets. The bottom figures show the same metrics for each algorithm in simulated *D. melanogaster* datasets: **D.** sensitivity, **E.** precision, and **F.** F1 score. Data from coverage levels of 1x, 2x, 3x, 4x, 5x, 10x, 20x, 30x, 40x, and 50x are displayed. Each algorithm is color-coded: LOCATE in red, xTea in green, TLDR in light blue, TrEMOLO in brown, TELR in grey, GraffiTE in pink, MEHunter in purple, PALMER in dark blue, TEMP2 in yellow, and MELT in orange. This color scheme is consistent throughout the study.

The sensitivity of LOCATE in detecting transposon insertions increases with sequencing coverage, rising from 17.27% at 1x coverage to 98.75% at 30x coverage, and nearly saturating at higher coverages with HiFi data (Fig. 2A; Table S2). Other algorithms follow similar trends but demonstrate lower sensitivity compared to LOCATE (Fig. 2A and Fig. S2A; Table S2; sensitivity drops from 98.08% to 42.33% at 30x coverage across three platforms). Long-read-based algorithms generally maintain stable sensitivity across the three platforms, except for xTea and TELR, which fail to detect most transposon insertions when using CLR reads with ∼10% error rates (Fig. S2A; top panel). Half of the long-read-based algorithms (LOCATE, xTea, TLDR, and TrEMOLO) are more sensitive than short-read-based methods (TEMP2 and MELT) in detecting transposon insertions using HiFi data (Fig. 2A; Table S2). LOCATE’s application of AutoML and post-filtering allows it to achieve 100% precision in detecting transposon insertions in simulated datasets from all three platforms, across nearly all coverage depths (Fig. 2B and Fig. S2B; Table S2). While five other algorithms (xTea, TLDR, PALMER, MEHunter, and TELR) also reach 100% precision with HiFi reads, none of them can reliably filter false positives in ONT and CLR reads (Fig. 2B and Fig. S2B; Table S2).

We further evaluated the F1 scores, which balance both sensitivity and precision, for detecting transposon insertions in the human simulated datasets (Fig. 2C and Fig. S2C; Table S2). LOCATE achieved the highest F1 score across all three platforms, demonstrating its ability to filter false positives from imperfect alignments while retaining most true positives (Fig. 2C and Fig. S2C). Among the ten benchmarked algorithms, TrEMOLO performs second best, followed by xTea, TLDR, PALMER, TEMP2, GraffiTE, MELT, TELR, and MEHunter when using simulated 30x human HiFi data (Fig. 2C).

The distribution of simulated human transposon insertions are based on previous findings, with Alu, a ∼280 bp element, contributing the majority (83%) of newly identified transposon insertions (12, 15). Since the length and class of insertions can influence detection performance, we next benchmarked LOCATE and other algorithms using simulated *D. melanogaster* datasets, which feature longer transposon insertions, including many LTR and DNA elements (Fig. 2D–F and Fig. S2D–F; see Methods). LOCATE again excels in sensitivity and precision across all platforms in detecting transposon insertions in *D. melanogaster* data (Fig. 2D–F and Fig. S2D–F). Interestingly, the performance gap between LOCATE and other long-read-based algorithms is more pronounced in simulated *D. melanogaster* datasets than in simulated human datasets, indicating that LOCATE offers more stable performance regardless of insertion length or class (Fig. 2D–F vs Fig. 2A–C; Fig. S2D–F vs Fig. S2A–C).

In addition to the highest sensitivity and precision, LOCATE can also accurately identify the breakpoints of transposon insertions; the average distances between LOCATE-predicted and simulated insertion breakpoints approach zero for both human and *D. melanogaster*, indicating near-perfect localization of insertion sites (Fig. S3).

In summary, LOCATE outperforms all other algorithms in identifying transposon insertions using simulated human and *D. melanogaster* datasets, showing the highest sensitivity and precision across different sequencing platforms, coverage levels, and insertion types.

### Assembling transposon insertion sequences in simulated datasets

Transposons often accumulate structural and single-nucleotide mutations during or shortly after replication (47–50). Consequently, accurately assembling transposon insertion sequences is essential for understanding their evolution and insertion mechanisms. Using simulated datasets, we evaluated the assembly accuracy of LOCATE, xTea, and TLDR, excluding other algorithms due to their lack of assembly functionality (TrEMOLO, GraffiTE, PALMER, TEMP2, and MELT) or low sensitivity (TELR and MEHunter). xTea is not applicable to *D. melanogaster* and not included into the benchmark in simulated *D. melanogaster* datasets. Among the tested algorithms, LOCATE consistently assembles insertion sequences with the fewest errors across all platforms (Fig. 3; Table S3).

**Figure 3.**
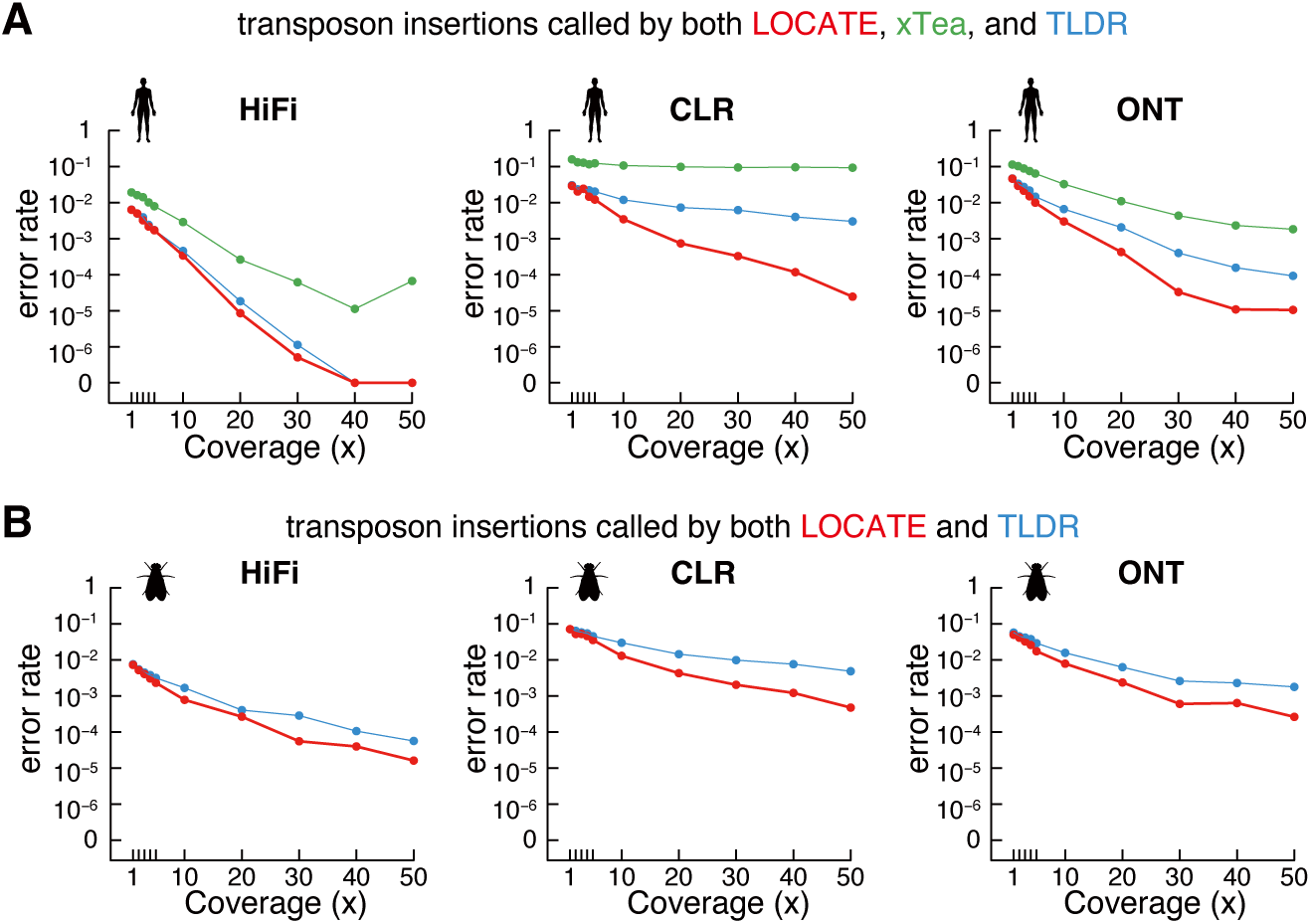
LOCATE assembles transposon insertion sequences with fewest errors in simulated human and *D. melanogaster* datasets. **A.** Line plots show the log-scaled error rate of transposon insertion sequence assemblies for LOCATE, xTea, and TLDR across three sequencing platforms in simulated human datasets. Only insertions detected by all three algorithms are included in the analysis. **B.** Same as A but for simulated *D. melanogaster* datasets.

Same as sensitivity, assembly accuracy was positively correlated with sequencing coverage across all platforms. At 1x coverage, the error rates of assembled sequences reflected the error rates of the respective sequencing platforms but decreased significantly with increasing coverage (Fig. 3; Table S3). When comparing transposon insertions identified by all benchmarked algorithms, LOCATE consistently yielded the lowest error rates across simulated human and *D. melanogaster* HiFi, CLR, and ONT datasets (Fig. 3). Using simulated 40x and 50x human HiFi data, LOCATE perfectly assembled insertion sequences (Fig. 3A; average error rate = 0). Although TLDR demonstrated accuracy comparable to LOCATE when using simulated high-coverage human HiFi data, both TLDR and xTea assembled insertion sequences with substantially higher error rates when analyzing simulated human CLR and ONT data (Fig. 3A). For 50x simulated CLR and ONT datasets, LOCATE maintained an average error rate of the level of 10⁻⁵, while TLDR and xTea exhibited error rates that were at least 8.1-fold higher than those of LOCATE (Fig. 3A). Notably, the assembly accuracy of LOCATE and TLDR is lower in simulated *D. melanogaster* datasets, likely due to their longer transposon insertions, than simulated human datasets (Fig. 3B).

In summary, LOCATE achieves error rates as low as zero using simulated 40x and 50x human HiFi data, 10^−5^ with simulated 50x human CLR and ONT data, and 10^−3^ to 10^−5^ in simulated 50x *D. melanogaster* datasets. This high precision makes LOCATE a robust tool for assembling and characterizing transposon insertions across diverse sequencing platforms and species.

### Identification of transposon insertions in real biological data

Real biological data present greater complexity than simulations, making it essential to evaluate the performance of LOCATE in practical scenarios. To this end, we applied LOCATE to the Genome In A Bottle (GIAB) dataset, which is widely regarded as the benchmark for structural variation detection (see Methods) (45). TELR and MEHunter showed low performance in simulated datasets and were excluded from the benchmark analysis (Fig. 2). We focused our assessment on LINE1, Alu, and SVA, as only three HERV-K insertions were detected by these algorithms. Across these datasets, LOCATE shows higher sensitivity and precision compared to the other seven algorithms (Fig. 4).

**Figure 4.**
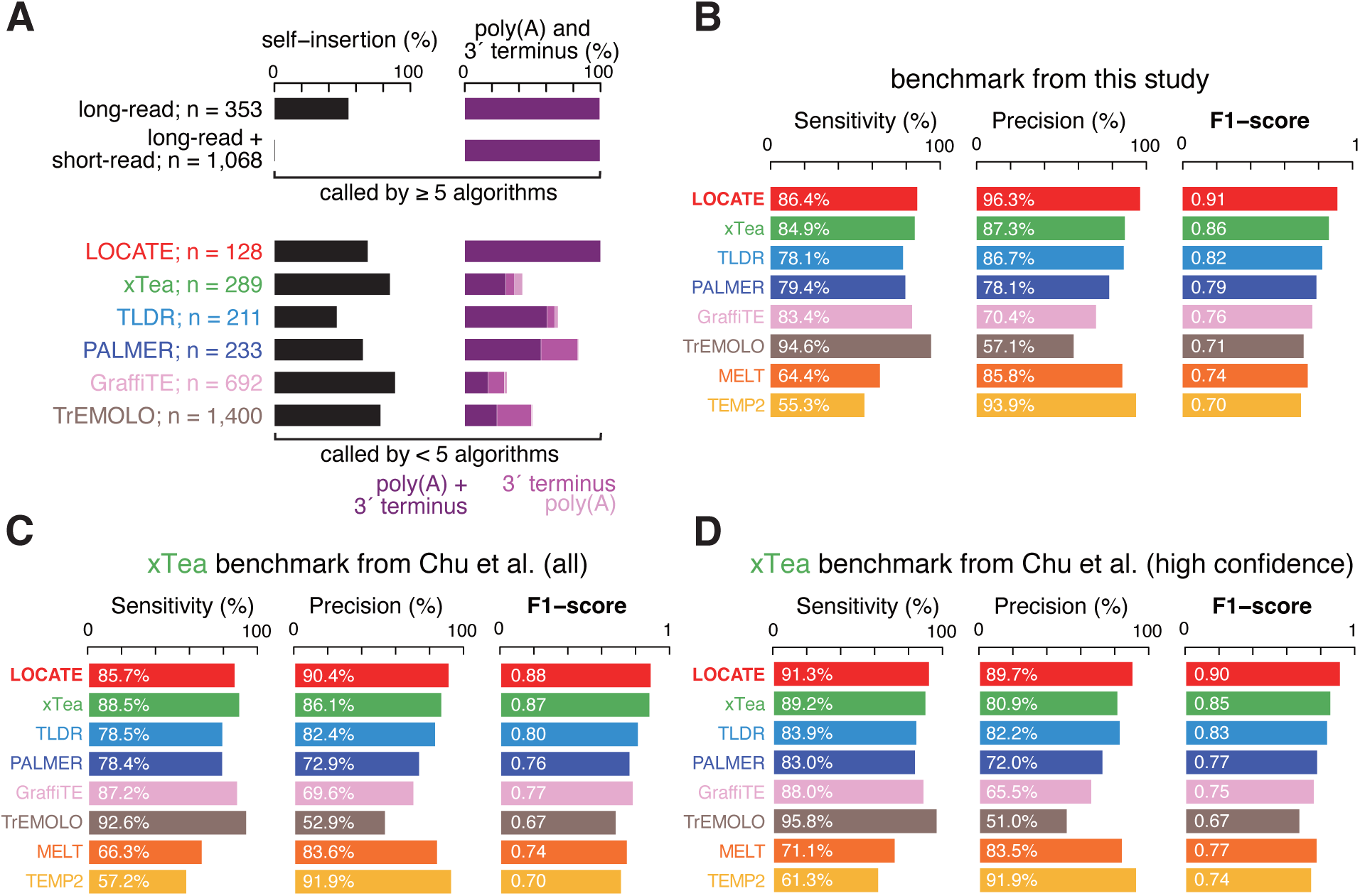
LOCATE outperforms other algorithms in detecting transposon insertions in GIAB datasets. **A.** Bar plots indicate the percentage of self-inserted transposon insertions (left) and transposon insertions with poly(A) and/or complete 3′ termini (right). Transposon insertions are divided into two groups: called by at least five algorithms (top two bars) and called by less than five algorithms (bottom six bars). Those called by less than five algorithms are further stratified by six long-read-based algorithms. Transposon insertions with both poly(A) and complete 3′ termini are colored in dark purple, while those containing only one of the two features are colored in purple and light purple. **B.** Sensitivity, precision, and F1 score for LOCATE, xTea, TLDR, PALMER, GraffiTE, TrEMOLO, TEMP2, and MELT in detecting transposon insertions in the GIAB HG002 HiFi dataset. Transposon insertions detected by at least five algorithms and those detected by less than five algorithms but passed manual examination are included in the benchmark from this study. **C–D.** Same as B, but assessed in the raw (**C**) and high confidence xTea benchmarks (**D**) from Chu et al.

Using HiFi sequencing data from HG002 (the son in the GIAB trio), LOCATE and other algorithms collectively identify 3,316 new transposon insertions (Table S4). Among these, 1,421 insertions were detected by more than five algorithms, 853 were identified by fewer than five but more than one algorithm, and 1,042 were unique to a single algorithm (Fig. S4A). Since different algorithms exhibit distinct error spectra, a stringent common set (insertions detected by at least five algorithms) is likely to minimize false positives. Supporting this, 99.51% of insertions in the common set contained complete 3′ ends and poly(A) tails, consistent with the insertion mechanisms of LINEs, SINEs, and retroposons (Fig. 4A; top panel). We further stratified the 1,421 common insertions into two groups based on whether they are called by short-read-based algorithms (TEMP2 and MELT). Interestingly, only six out of the 1,068 common insertions called by both long-read-based and short-read-based algorithms are self-insertions, significantly less than those only called by long-read-based algorithms (193 out of 353; Chi-squared *p*-value < 2.2 × 10^−16^), suggesting the advantage of long-read in identifying self-insertions (Fig. 4A; top panel).

We then carefully inspected those transposon insertions called by each of the six long-read-based algorithms but supported by less than 3 additional algorithms (Fig. 4A; bottom panel; n = 128 to 1,400 for these algorithms). A significant portion of these transposon insertions are self-insertions (46.0% to 88.9%), that represent the most complex cases in which many false positives exist due to alignment errors, incomplete reference assemblies, or other structural variations within pre-existing transposons (Fig. 4A; bottom panel). A bona-fide LINE1/Alu/SVA insertion will have complete 3′ termini and extra poly(A) tail (34, 35). However, many of these transposon insertions called by the other five long-read-based algorithms don’t have a complete 3′ terminus or a poly(A) tail (Fig. 4A; bottom panel; 39.3% for TLDR to 82.8% for GraffiTE), suggesting false positives widely exist in these algorithms.

To systematically evaluate LOCATE and other algorithms, we constructed a gold-standard dataset comprising the common insertions and the other insertions with manual examination in the genome browser (Fig. S4B and S5; Table S5; see Methods). We manually examined 792 transposon insertions called by less than five algorithms with complete 3′ termini. In the 792 transposon insertions, 56 were designated as false positives as they were from locus with extremely high coverage (Fig. S5A; >5 copy number). We further filtered 358 transposon insertions as the majority of the insertion sequences were not from transposons, and the non-transposon part were not from transduction (Fig. S5B). 67 transposon insertions are only supported by multi-mappers (reads mapped to multiple loci with map quality equal to zero). We found these multi-mappers often result in extra haplotypes around the insertion breakpoints. For instance, a transposon insertion called by TLDR is only supported by multi-mappers; while the unique-mappers suggest two haplotypes around the breakpoint, multi-mappers fall into neither of the haplotypes (Fig. S5C). Thus we designated the 67 transposon insertions as false positives. Finally, 311 transposon insertions passed our manual examination, and together with the common set, a gold-standard set with 1,732 insertions were used to evaluate the performance of LOCATE and other algorithms (Fig. S5D as an example).

As shown in Fig. 4B, LOCATE achieved the highest performance among all algorithms, with a F1 score of 0.91 compared to 0.69–0.86 for other algorithms (McNemar’s test *p*-values < 2.2 × 10^−16^). Among all algorithms, LOCATE recalls the second most true positive transposon insertions (sensitivity = 86.4%) while keeps the highest precision (96.3%) (Fig. 4B). While the other long-read based algorithms (xTea, TLDR, PALMER, GraffiTE, and TrEMOLO) show comparable sensitivities (78.1%–94.6%), none of them can reach precision as high as LOCATE (57.1%–87.3% vs 96.3%; 2.43– to 10.59–fold false discovery rate decreasing) (Fig. 4B). Short-read based algorithms (MELT and TEMP2) avoid identifying self-insertions and keeps high precision (85.8%– 93.9%), but they are less sensitive compared to long-read based algorithms (55.3%–64.4%) (Fig. 4B).

We further employed other benchmarks, which are used by xTea, as a supplement for evaluating LOCATE and other algorithms (15). The xTea raw benchmark, which consists of 1,642 transposon insertions, is based on carefully benchmarked insertion calls from GIAB and a haplotype-resolved assembly, followed by using repeatMasker to annotate transposon insertions and manual confirmation for poly(A) tail and TSD (15, 45, 51). The xTea team generated a highly confident benchmark, requiring manual confirmation for the full TPRT signature, yielding 1,531 transposon insertions (15). Most transposon insertions from the two xTea benchmarks are supported by multiple algorithms, especially the high confidence benchmark (Fig. S4B). Again, LOCATE showed the highest performance in both the raw and the high confidence xTea benchmarks (Fig. 4C–D; F1 = 0.90 for LOCATE vs 0.67–0.85 for other algorithms in the high confidence benchmark; McNemar’s test *p*-values < 0.04; F1 = 0.88 for LOCATE vs 0.67–0.87 for other algorithms in the low confidence benchmark; McNemar’s test *p*-values < 1.9 × 10^−21^). Assessed on our manual examined benchmark and two benchmarks from the xTea study, LOCATE shows significantly higher performance compared to other algorithms in identifying new transposon insertions.

### Alu prefers two specific hotspots for self-insertions

The enhanced sensitivity and accuracy of transposon insertion detection achieved by LOCATE, particularly in identifying self-insertions, allow for a detailed study of insertion site preferences and transposition mechanisms. Applying LOCATE to the GIAB HG002 dataset, we identified 183 out of 1,304 Alu insertions that occurred within Alu elements in the reference genome. These Alu self-insertions show a significant preference at two distinct hotspots in its consensus sequence (Fig. 5A).

**Figure 5.**
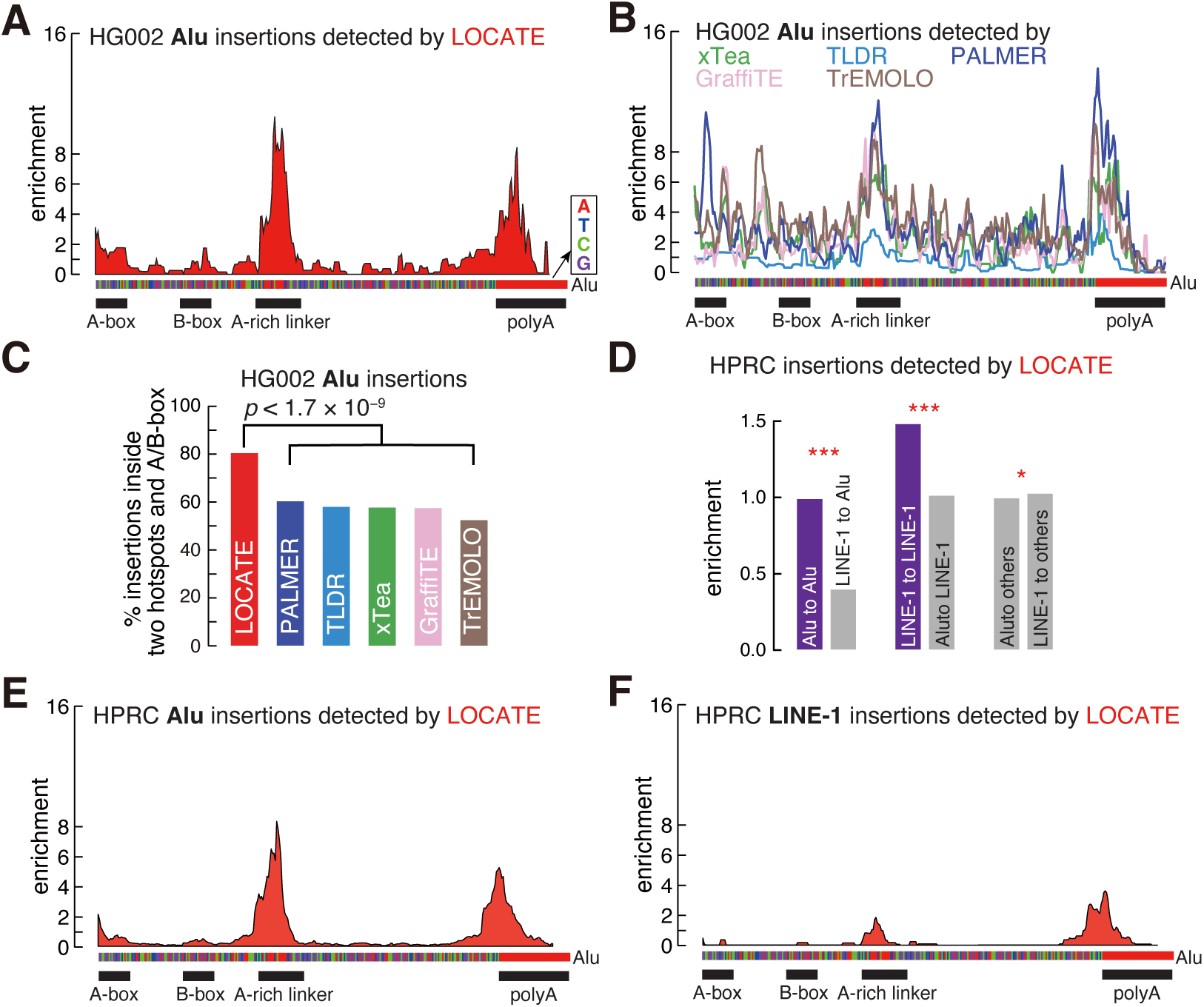
Alu prefers self-insertion in two hotspots. **A.** Bar plots present the enrichment of LOCATE-detected HG002 Alu self-insertions at each nucleotide of the Alu consensus sequence. The Alu consensus sequence is shown at the bottom, with the A-box, B-box, A-rich linker, and poly(A) tail regions marked. **B.** Same as A but for HG002 Alu self-insertions detected by the other five long-read-based algorithms. **C.** Percentage of HG002 Alu self-insertions detected by LOCATE and other algorithms from Alu regions inside the two hotspots, A-box, and B-box. **D**. Enrichment over expectation of LOCATE-detected HPRC Alu and LINE1 insertions at the pre-existing Alu copies, pre-existing LINE1 copies, and other genomic regions. Self-insertions are colored in purple. *: *p* < 0.05, ***: *p* < 0.001. **E.** Same as A but for HPRC Alu self-insertions detected by LOCATE. **F.** Same as A but for HPRC LINE1 insertions at Alu detected by LOCATE.

The most prominent hotspot, showing up to 10–fold enrichment over expectation, is located at the A-rich linker that separates the two 7SL-like dimers of the Alu element (Fig. 5A). Alu elements, which utilize TPRT mechanism, favor AT-rich sites for cleavage, priming, and pairing with their poly(A) tails during reverse transcription and genomic integration (Fig. S6A) (34, 35). This aligns with our observation that the A-rich linker is the most preferred site for Alu self-insertion. Similarly, the poly(A) tail at the 3′ end of Alu is the second most enriched hotspot, showing up to 8–fold enrichment over expectation (Fig. 5A). In addition to the two hotspots, we also observed two small peaks with modest enrichment at the Alu A-box and B-box. The small peaks of Alu self-insertions in the A-box and B-box are consistent with studies suggesting that non-LTR retrotransposon insertions slightly favor open chromatin regions, which are more accessible to integration machinery (52–55).

We compared the results from LOCATE with those from other long-read-based algorithms. While Alu self-insertions detected by other algorithms also enrich the two hotspots, a higher percentage of these self-insertions are in non-hotspot surrounding regions, indicating they might carry more false positives (Fig. 5B; Fig. S6B). Specifically, 81% of the LOCATE identified Alu self-insertions are located in the two hotspots, A-box and B-box, significantly higher than those identified by other algorithms (52% to 60%; Chi-squared test *p*-value < 1.7 × 10^−9^). The comparison between LOCATE and other algorithms reinforces LOCATE’s superior accuracy in identifying self-insertions.

To explore more of transposon insertions at Alu, we applied LOCATE to the 51 additional human individuals from the Human Pangenome Reference Consortium (HPRC) (Table S6) (56), identifying 10,751 Alu, 1,728 LINE1, and 268 SVA germline insertions. The analysis in HPRC datasets replicated our findings using the GIAB HG002 dataset (Fig. 5D–E). 1,202 out of the 10,751 Alu are self-insertions, and these Alu self-insertions are enriched in the two hotspots: A-rich linker and poly(A) tail (Fig. 5E). Interestingly, most Alu self-insertions at the two hotspots are in the sense strand of the pre-existing Alu, consistent with the mechanism that the 3′ end of Alu transcripts targets poly(T) in the negative strand of pre-existing Alu hotspots, reverse transcribe, and produce new insertions with the same/sense strand of the pre-existing Alu (Fig. S6C).

Including more individuals from HPRC allowed us to identify more LINE1 insertions and investigate their enrichment at pre-existing Alu insertions. LINE1 uses the same mechanism, endonuclease, and reverse-transcriptase for reverse transcription and integration as Alu. Recapitulating previous findings, both Alu and LINE1 favors the same sequence motif AA/TTTT for insertion, with “/” as the insertion site (Fig. S6A). LINE1 insertions are enriched at the same two hotspots at the Alu consensus (Fig. 5F). LINE1 also favors insertion at the sense strand of the two Alu hotspots (Fig. S6D). However, the enrichment of LINE1 insertions at the two Alu hotspots are ∼2–fold weaker than Alu insertions (Fig. 5F vs. Fig. 5E). Overall, Alu is more likely to integrate into a pre-existing Alu than LINE1 (Fig. 5D; Chi-squared test, *p*-value < 2.2 × 10^−16^), while a LINE1 insertion is more likely to occur in pre-existing LINE copies than an Alu insertion (Fig. 5D; Chi-squared test, *p*-value = 2.1 × 10^−8^). Previous studies have shown many DNA transposons like to insert into nearby genomic regions, forming a phenomenon called “local hopping” (57–60). Another study found LINE1 and Alu are separately clustered and preferentially localized to different chromosome compartments (61). Thus, we speculate that Alu and LINE1, like DNA transposons, have the same “local hopping” phenomenon, that after transcription, their RNAs can immediately enter the transposition machinery at nearby regions, leading to the higher rate of self-insertions than non-self-insertions.

By applying LOCATE to GIAB and HPRC datasets, we identified the Alu A-rich linker and poly(A) tail as two hotspots for Alu and LINE1 insertions. We also showed a higher frequency of self-insertions than non-self-insertions for Alu and LINE1. These results suggest a feedforward mechanism for human transposon replication, by which pre-existing insertions provide favorable sites for subsequent integration of its own transcripts.

## Discussion

Transposon activity is a double-edged sword in genome biology. On one hand, transposons drive population diversity and genome evolution; on the other hand, they can disrupt gene expressions, compromise genome stability, and cause diseases (2–9). While short-read algorithms for detecting transposon insertions are limited by read length and unable to resolve repetitive regions, long-read sequencing offers a promising avenue for overcoming these challenges. However, existing algorithms have struggled to meet this challenge. In response, we developed LOCATE, a novel algorithm that outperforms others in identifying transposon insertions and assembling their sequences across diverse species and sequencing platforms.

Many long-read-based algorithms, such as TrEMOLO, GraffiTE, and MEHunter, depend on structural variation callsets generated by tools like Sniffles/Sniffles2 or genome assemblies, followed by distinguishing transposon insertions from other structural variation (13, 14, 17, 62, 63). This approach is constrained by the quality of the initial structural variation callset and the secondary discrimination strategy. LOCATE avoids these pitfalls by implementing an alignment-based method that directly searches for transposon insertion candidates in reads aligned to both the genome and transposon sequences. The other algorithms also use the same strategy for identifying transposon insertions, but LOCATE is unique from its three core modules: machine learning-based filtering of alignment artifacts, precise assembly and characterization of insertion sequences, and family-specific post-filtering based on insertion characteristics. Furthermore, LOCATE maintains computational efficiency, outperforming most other algorithms in runtime (Fig. S7; 3–fold to 679–fold faster than other algorithms except MEHunter).

One of the most significant challenges in transposon detection is accurately identifying self-insertions (10, 15, 18, 52). The repetitive nature of transposons made these self-insertions vulnerable to artifacts from many sources, such as inaccurate alignments and other structure variations. Short-read-based methods typically exclude such insertions to limit false positives (10, 11), while long-read algorithms have not yet fully addressed these challenges. LOCATE excels in this area by combining machine-learning models with post-filtering based on key insertion features. While its improvements over algorithms like xTea and TLDR might appear incremental (Figs. 2 and 4), LOCATE’s capability to resolve self-insertions enables detailed genome-wide analyses. For instance, applying LOCATE to GIAB and HPRC datasets suggests Alu elements preferentially insert into two hotspots within themselves, highlighting a potential "feedforward" mechanism facilitating their genome integration.

Despite its strengths, LOCATE has some limitations. Its sensitivity in extremely repetitive regions, such as telomeres and centromeres, is constrained by the need for ultra-long reads. Additionally, LOCATE’s reliance on post-filtering based on general insertion features may limit its ability to detect transposons with atypical integration mechanisms. For example, R2 transposons, which also use TPRT for reverse transcription and integration but lack poly(A) tails in their insertions, might escape detection under these criteria (64–66). Thus, users should be aware of those insertions without passing the post-filtering step. Furthermore, the machine-learning models for filtering alignment artifacts should be species-specific. While pre-trained models are available for humans, *D. melanogaster*, and mice, users working on other species need to invest additional resources and time in training these models.

## Acknowledgements

We would like to thank the members of Weng laboratories for their critical comments and discussion on the manuscript.

## Author Contributions

T.Y. and Z.W. conceived the project. Z.H. and B.X. developed LOCATE. B.X. and Z.H. benchmarked LOCATE and other algorithms. B.X. and Z.H. analyzed the preference of Alu and LINE1 insertion sites. T.Y. wrote the manuscript, with the input and suggestions from all authors.

## Data Availability

LOCATE is freely available at GitHub: https://github.com/red-t/LOCATE. HiFi sequencing data for HG002 was downloaded from GIAB: https://ftp.ncbi.nlm.nih.gov/ReferenceSamples/giab/data/AshkenazimTrio/HG002_NA24385_son/. HiFi sequencing data for 51 additional human individuals were downloaded from HPRC Github page: https://github.com/human-pangenomics/HPP_Year1_Data_Freeze_v1.0.

## Supplementary figure legends

**Figure S1 (Related to Fig. 1).**
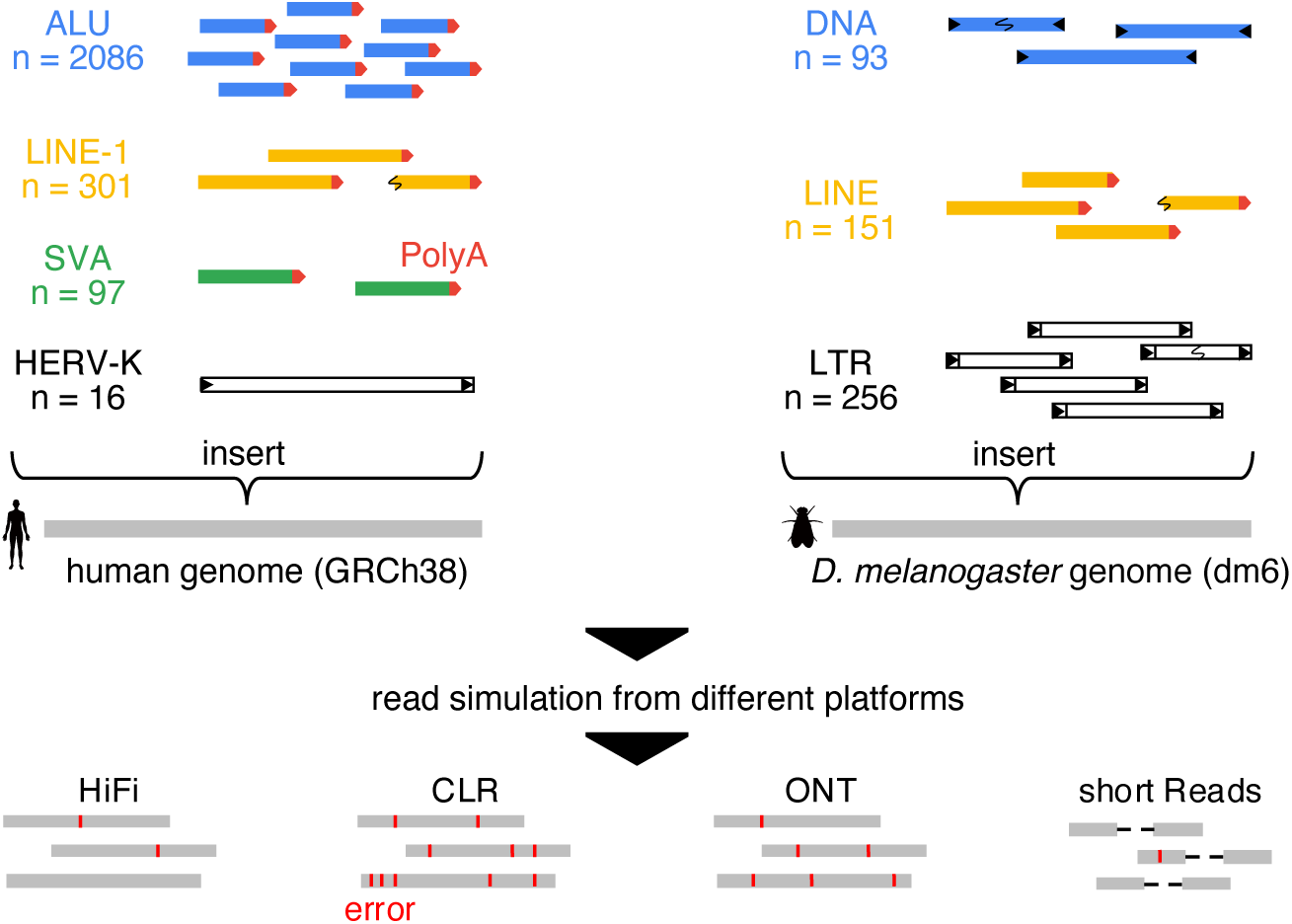
Schematic of data simulation for individual human genomes and *D. melanogaster* population genomes.

**Figure S2 (Related to Fig. 2).**
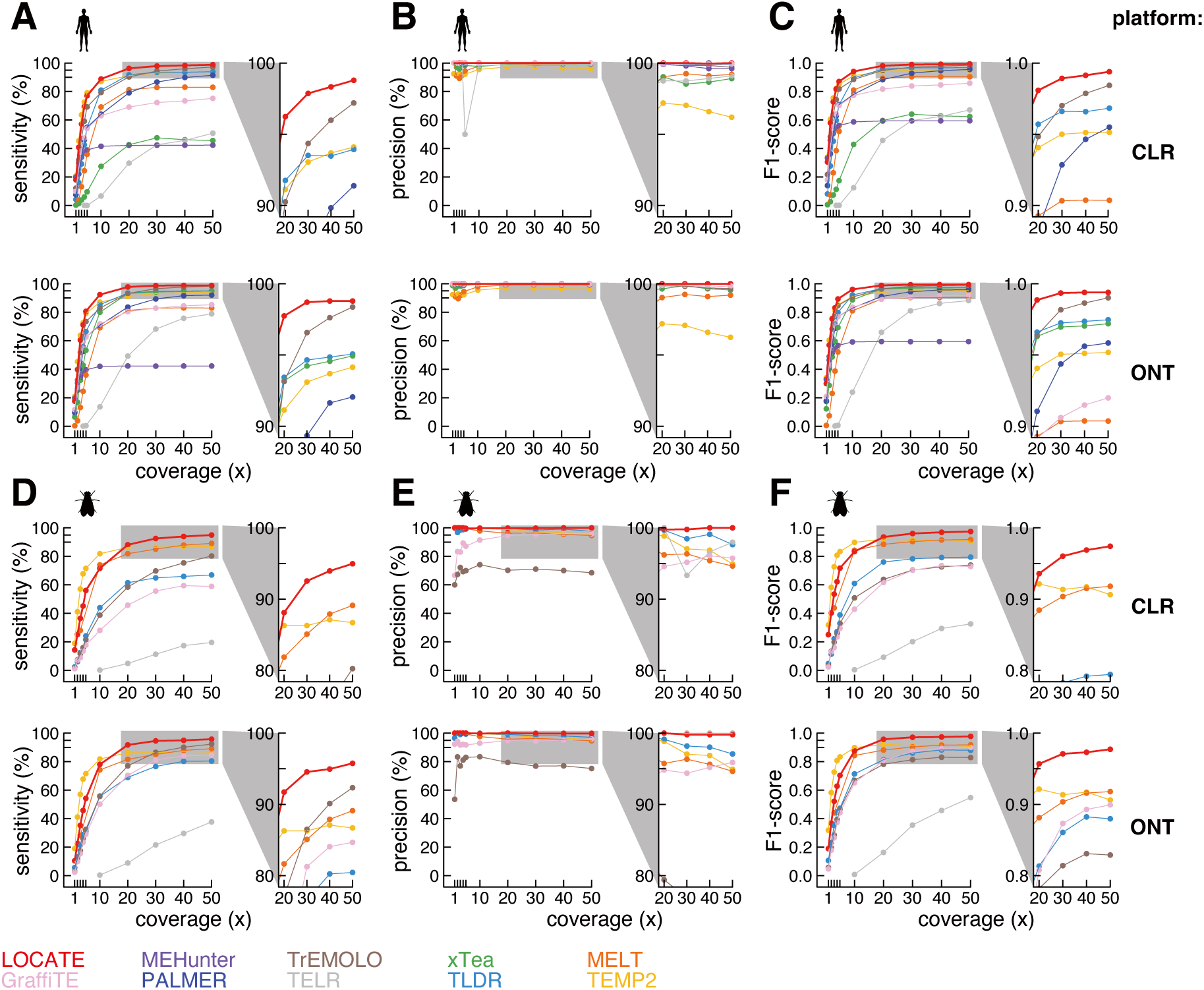
LOCATE outperforms other algorithms in detecting transposon insertions in simulated human and *D. melanogaster* datasets. A–C. Same as Fig. 2A–C but in the other two long-read sequencing platforms: Pacbio CLR and ONT. **D–F.** Same as A-C but benchmarked in simulated *D. melanogaster* datasets.

**Figure S3 (Related to Fig. 2).**
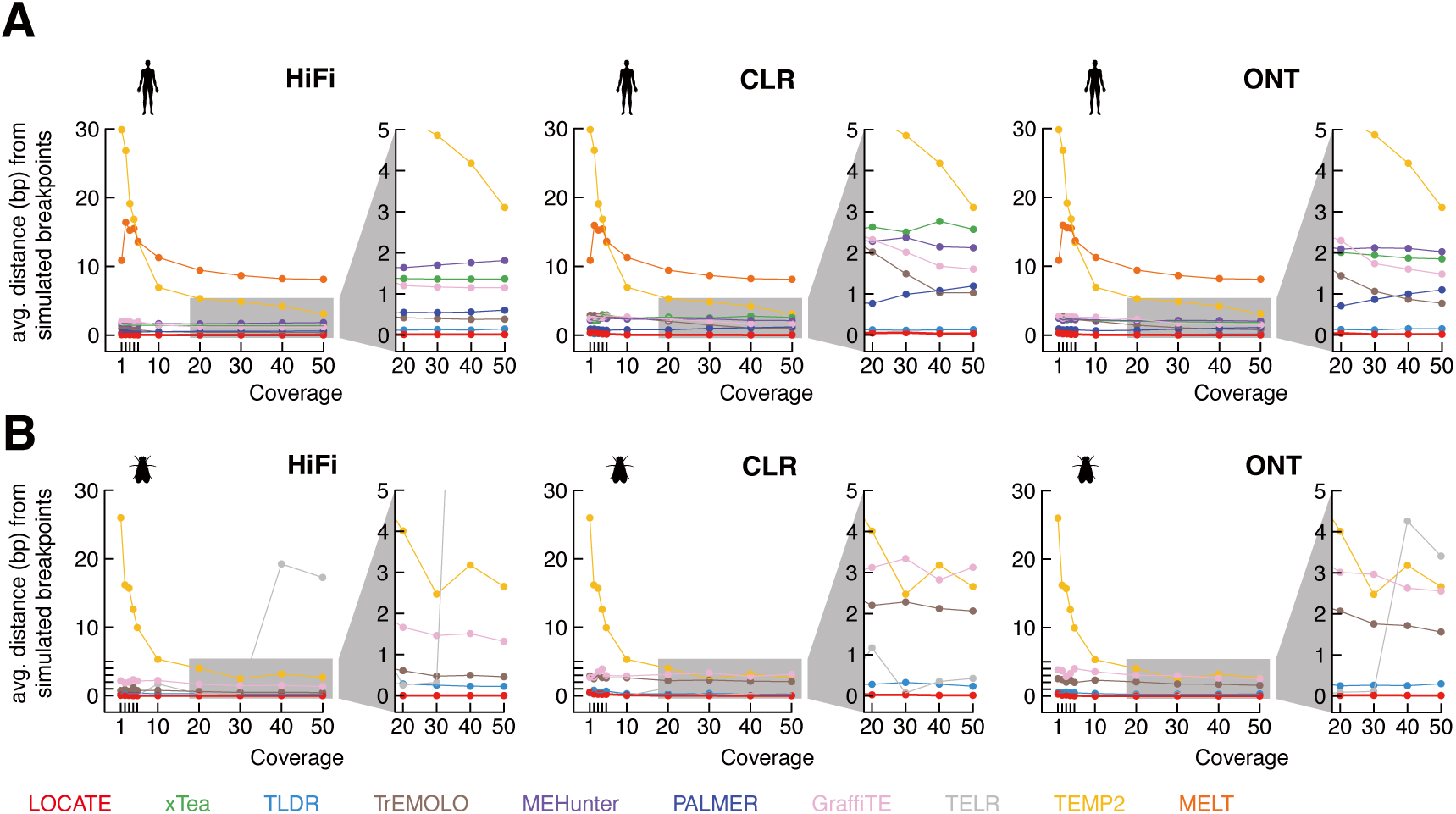
LOCATE performs best in identifying transposon insertion breakpoints. **A.** Average distance between detected and simulated insertion breakpoints in simulated human datasets across three platforms. **B.** Same as A but for simulated *D. melanogaster* datasets.

**Figure S4 (Related to Fig. 4).**
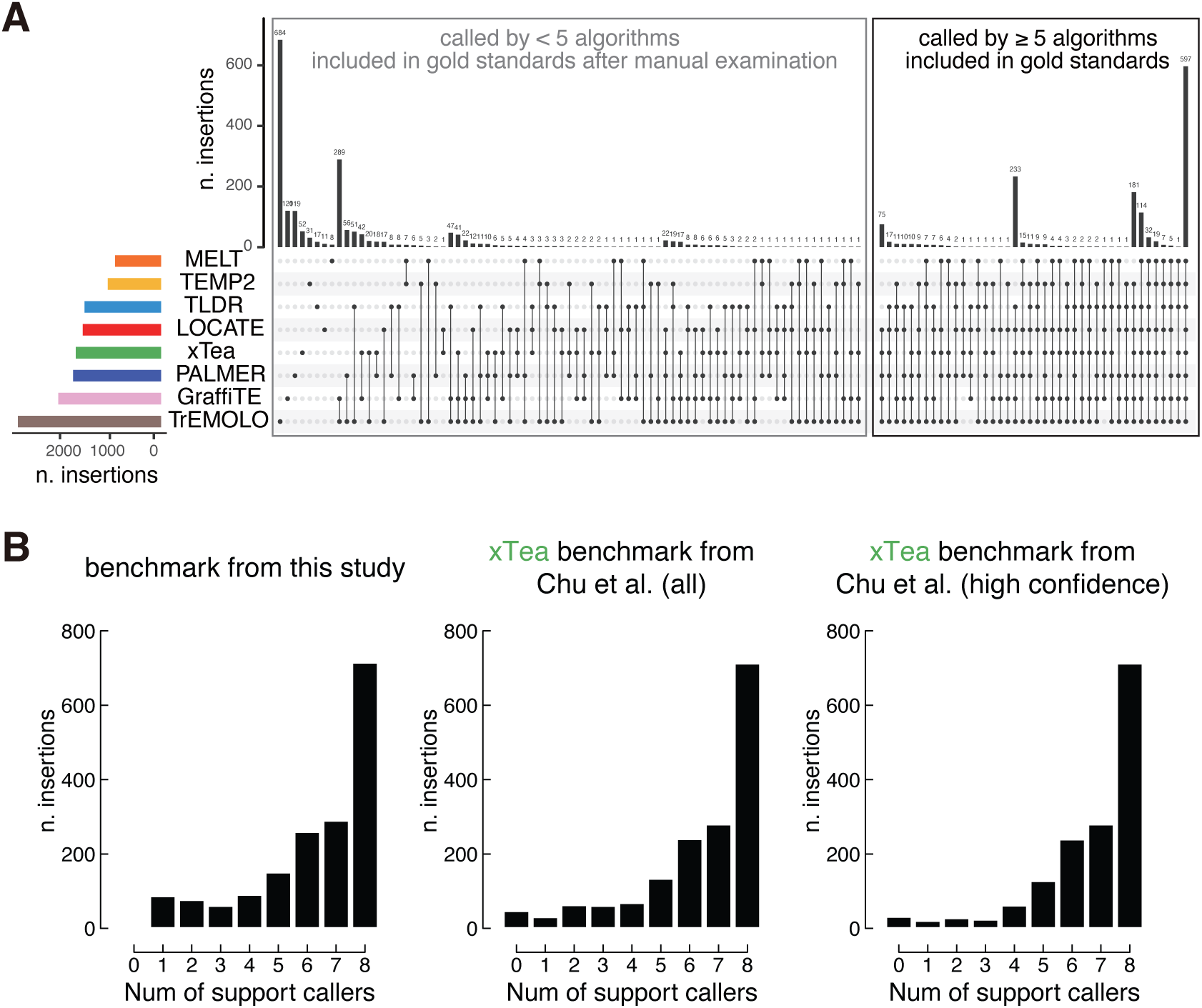
Overview of transposon insertions detected by LOCATE and other algorithms using the GIAB dataset. **A.** An upset plot shows the overlap of transposon insertions detected by LOCATE and other algorithms in the GIAB HG002 HiFi dataset. Transposon insertions called by at least five algorithms are marked in the black box and directly included into the gold standards, while those called by less than five algorithms are marked in the gray box and included into the gold standards after passing manual examination. **B.** Histograms indicate how many algorithms can detect the transposon insertions defined in our benchmark, xTea raw benchmark, and xTea high confidence benchmark.

**Figure S5 (Related to Fig. 4).**
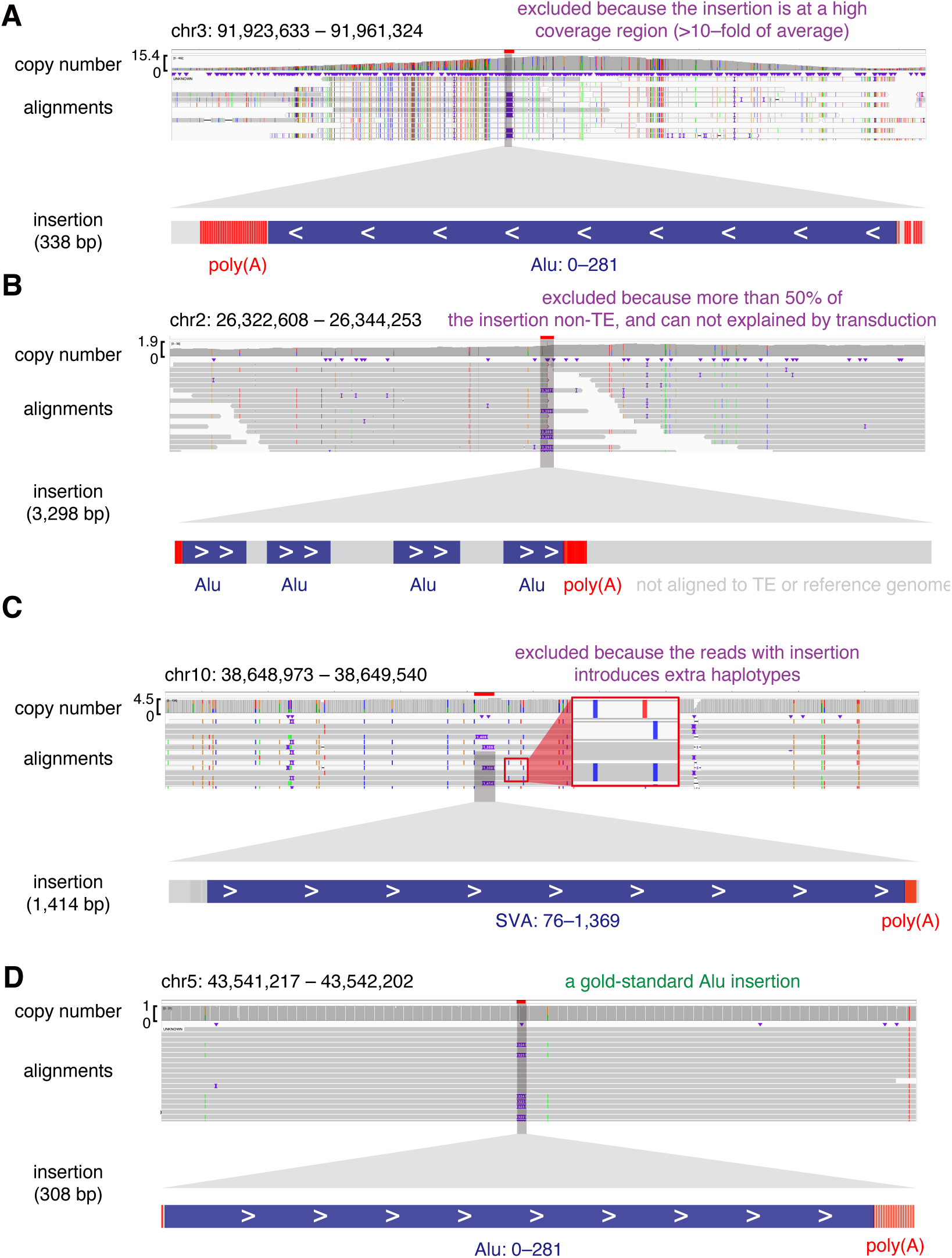
Representative examples of manual examined transposon insertions. **A.** A false positive transposon insertion located at a genomic region with extremely high coverage. This false positive likely originates from incomplete genome assembly. **B.** A false positive transposon insertion without generic transposon structure. Its inserted sequence contains four Alus. While no transduction is observed, this insertion is likely not derived from transposition. **C.** A false positive transposon insertion only supported by multi-mappers. While the unique-mappers suggest for two haplotypes upstream this transposon insertions, the multi-mappers align with neither of the two haplotypes, suggesting inaccurate alignment. **D.** A true positive Alu insertion, which has a poly(A) tail and complete 3′ terminus, is supported by uniquely mapped reads, and falls in a region with normal coverage.

**Figure S6 (Related to Fig. 5).**
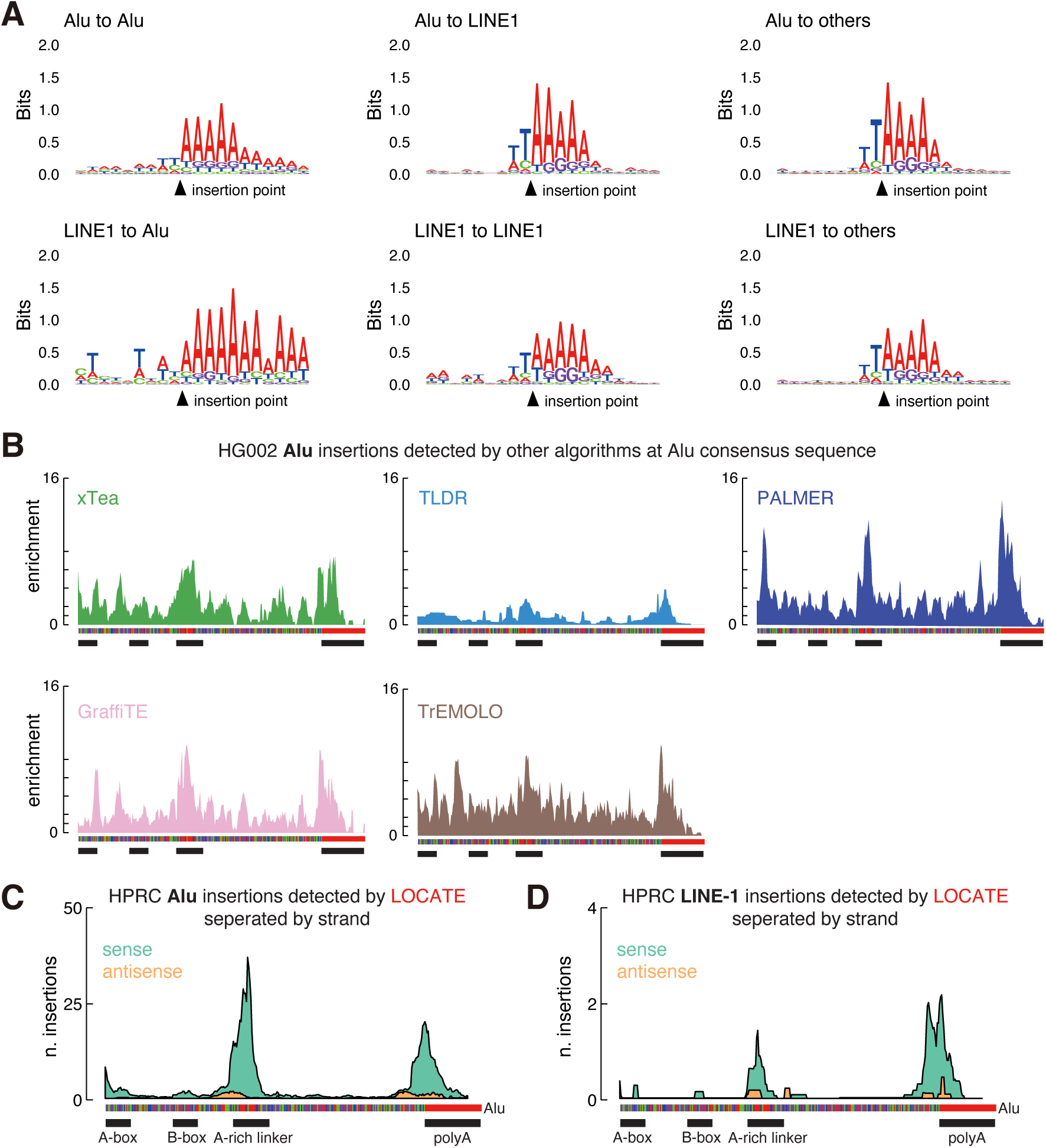
Alu and LINE1 favor inserting into AT-riched regions and the sense strand of pre-existing Alus. **A.** Sequence logos around the insertion points (5′ breakpoints) of Alu and LINE1 insertions at pre-existing Alu copies, pre-existing LINE1 copies, or other genomic regions. **B.** Similar to Fig. 5E. The enrichment of LOCATE-detected HPRC Alu self-insertions at each nucleotide of the Alu consensus sequence. Insertions in sense and antisense Alu strand are colored in cyan and orange respectively. **C.** Same as panel A, but for LINE1 insertions at pre-existing Alus.

**Figure S7 (Related to Discussion).**
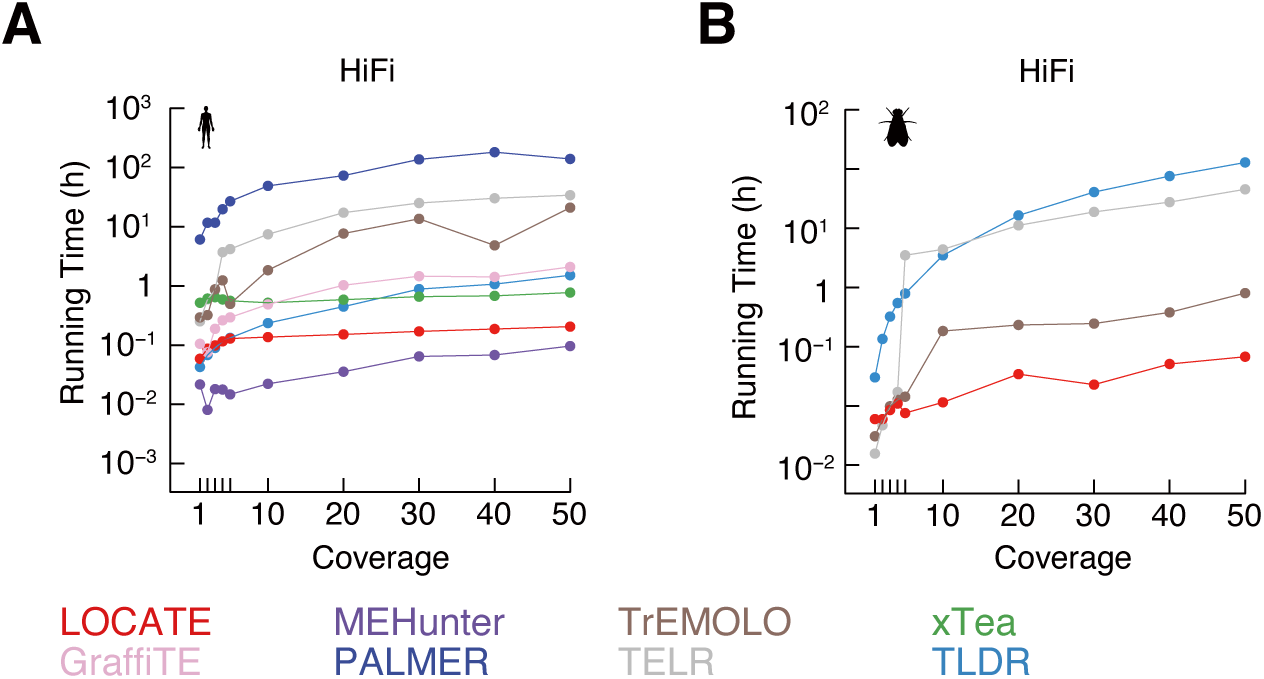
LOCATE requires less runtime for detecting transposon insertions than most of the other algorithms. Runtime for identifying transposon insertions in simulated human (**A**) and *D. melanogaster* (**B**) HiFi datasets. The eight long-read-based algorithms are shown. Sequencing data from 1x to 50x coverages are benchmarked.

## Supplementary table legends

**Table S1. Detailed descriptions of simulated transposon insertions in human and *D. melanogaster* genomes.**

**Table S2. Performance metrics (sensitivity, precision, and F1 scores) for LOCATE and other transposon detection algorithms across simulated datasets.**

**Table S3. Error rates in transposon sequence assembly for LOCATE and other algorithms in simulated datasets.**

**Table S4. Transposon insertions detected in the GIAB HG002 HiFi dataset by LOCATE and other algorithms.**

**Table S5. Gold-standard transposon insertions used for benchmarking in this study, derived from the GIAB HG002 HiFi dataset.**

**Table S6. Transposon insertions detected in the 51 additional human individuals from HPRC by LOCATE.**

**Table S7. Detailed information of the 19 features used in AutoML model training for LOCATE.**

